# Accumulating waves of random mutations before fixation

**DOI:** 10.1101/2022.06.04.494827

**Authors:** Marius Moeller, Benjamin Werner, Weini Huang

## Abstract

Mutations provide variation for evolution to emerge. A quantitative analysis of how mutations arising in single individuals expand and possibly fixate in a population is essential for studying evolutionary processes. While it is intuitive to expect that a continuous influx of mutations will lead to a continuous flow of mutations fixating in a stable constant population, joint fixation of multiple mutations occur frequently in stochastic simulations even under neutral selection. We quantitatively measure and analyse the distribution of joint fixation events of neutral mutations in constant populations. We propose a new concept, the mutation “waves”, where multiple mutations reach given frequencies simultaneously. We show that all but the lowest frequencies of the variant allele frequency are dominated by single mutation “waves” that approximately follow an exponential distribution both in their size and the time between two “waves”. Consequently, large swaths of empty frequencies are observed in the variant allele frequency distributions, with a few frequencies having numbers of mutations far in excess of the expected average values. We further show that the discrete mutation waves average out to a continuous distribution named as the Wave Frequency Distribution (WFD), the shape of which is predictable based on few model parameters.

## 1 Introduction

Understanding the occurrence, expansion and fixation of spontaneous mutations has been an important topic in evolutionary biology [50]. Early studies involving small genomes like mitochondrial DNA [28, 51, 56] reported mostly sequential fixation of single mutations and rarely the fixation of multiple mutations [40]. With the development of new sequencing technologies, simultaneous fixation of multiple mutations has been observed more frequently, especially in larger genomes [12, 14, 27, 44, 61].

A number of hypotheses were proposed to explain the joint fixation of multiple mutations. For example, in the absence of crossovers or recombination such as in asexually reproducing cells, clonal interference [22, 25, 30, 33, 35] delays fixation and increases mutational loads [6, 8, 13]. Negative frequency-dependent selection (NFDS) is the situation where the fitness of mutations decrease as they reach high frequencies in a population [17, 24, 43, 52], which has been seen in many natural populations [53, 54]. Both clonal interference and negative frequency dependent selection result in the present of multiple sub-populations for an extended period of time. This allows many mutations to first accumulate in a smaller sub-population. Eventually, a strongly positively selected mutation may emerge in one of the sub-populations and quickly drives the expansion of this sub-population, which leads to its fixation in the entire population together with all mutations that accumulated in that sub-population. [38, 39, 44]. Theoretically, de Sousa et al. [47] have shown the prevalence of multiple mutations fixating simultaneously in simulations of Fisher’s Geometric Model. This model assumes mutation affecting fitness through the change in multiple traits, represented by a Gaussian distribution centered around zero. Each individual thus takes a position in a multidimensional fitness landscape, with the entire population being a cloud of these positions, moving towards local optima. This leads to frequent clonal interference and thus joint fixation of multiple mutations.

Other related hypotheses include the Staircase model [16, 18, 20, 29, 55], branching models [21, 46, 58] and the appearance of mutators [14, 42] with beneficial mutations. In the Staircase model, only specific beneficial mutations are tracked, and each mutation is assumed to have the same small selective advantage, akin to the population going up a staircase with each mutation. Simultaneous fixation of mutations has been found in this model [20, 29]. In branching models, cell populations “branch” into different sub-populations with different mutations, each of which can grow, and inter-competition between the sub-populations is often limited. There is a variety of different models that all are considered branching processes, but in one of the most well-known variants, the Wright-Fisher Process [1, 2], simultaneous fixations has been found in the context of jointly beneficial mutations, i.e. mutations that only have a positive effect if held together [19]. Lastly, the presence of mutator lineages that directly increase mutation rates. These mutators are likely to dominate when a colony is introduced into a new environment, where there is ample opportunity for new mutations to be beneficial [12]. Ultimately, once a strongly selected mutation emerges by chance, multiple mutations hitchhike with the mutator [14, 42].

While all the aforementioned hypothesis or models require strong selection [14, 38, 39, 42, 44, 47], is strong selection a necessary condition for simultaneous fixation of multiple mutations? Can neutral mutations jointly fixate in a population, especially under clonal reproduction where mutations on the same chromosome are linked? There are some related work on the accumulation of mutations in the “Muller’s Ratchet” models [6, 8–11, 15, 26, 57], where mutations with a very small but negative fitness effect are accumulated ultimately in a finite population despite being selected against long-term. In particular, Higgs and Woodcock [11] found extensive joint fixation of mutations due to the structure of the genealogical trees. Looking backward in a genealogical tree at the moment when a single mutation fixates, one will find a common ancestor among all currently alive individuals. All mutations present in this ancestor that have not fixated beforehand will fixate at once. This theory in principle is also valid to explain the joint fixation of neutral mutations independent of “Muller’s Ratchet” models. Measures such as the average mutation accumulation rate [8, 9], mean time to extinction [10] and diversity [15] have been addressed in these models. Despite of having these interesting results, quantitative understanding of joint fixation events are still missing, including e.g. the number of mutations that fixate together and the average time between two sequential fixation events. Here, we first present expressions for these two quantities. We extend our analyse by looking at the dynamics of mutation “waves” before fixation. Finally, we discuss how and why quantifying those detailed dynamics of fixation events and mutation “waves” helps to understand the variant allele frequency (VAF) distribution.

As we are interested in the joint fixation of random mutations in a stable population in the absence of strongly selected sub-populations, we use a minimal birth-death Moran process with neutral mutations and a constant mutation rate [4, 23, 29, 31, 34, 41]. Our stochastic simulations show that simultaneous fixation of multiple neutral mutations is not only possible but the norm, the distributions of which can be estimated by our analytical approximations. We then introduce a more general concept, we call “accumulating waves”, that describes how mutations jointly move towards high frequencies. This concepts applies to large parts of the VAF. It allows us to predict, e.g. the average time without any mutation present at any frequency as well as the amplitude and general shape of the mutation wave distribution.

Analytical solution of the VAF distribution both in growing and constant populations are usually continuous averages over stochastic fluctuations [3, 31, 37, 48]. However, in reality, the VAF distribution is highly non-continuous. Many especially high frequencies are entirely void of mutations, and the VAF distribution gets a bi-modal character. We show how the average behaviour of the VAF emerges from an effectively bimodal process with many empty frequencies and occasional waves of mutations. We predict the mean size of these waves, the time these waves are absent and show that they are approximately exponentially distributed for high frequencies. Importantly, this indicates a VAF distribution exceeding the expected average by large margins in real sequencing data is not necessarily an aberration in need of an explanation, but instead can be a regular occurrence even in the absence of selection.

## 2 Model Description

### 2.1 A birth-death Moran model with random neutral mutations

We model the accumulation of mutations in a constant population of size *N* utilising a birth-death Moran process [4]. Individuals reproduce and die at a rate *r* per individual per time unit. Reproduction or death events are happening according to an exponential distribution, which means that in a sufficiently small period of time 1 >> Δ*t*, every individual has a chance of *r*Δ*t* to reproduce. In a Moran process, reproduction and death happen simultaneously, with one individual producing two offspring to replace itself and a randomly chosen individual.

We model the dynamics in the context of an infinite allele model, where the chance to exactly hit a previously mutated location with a new mutation is zero because the genome size is large [11, 47, 48]. This implies a binomial distribution for the number of novel mutations across offsprings, which is well approximated by a Poisson distribution with the mutation rate *µ* as its mean [32]. For simplicity, we assume a constant mutation rate in our model. The mutation rate *µ* for multicellular organisms is generally thought to be in the order of 10^*−*9^ - 10^*−*7^ per base pair per reproduction, depending on the concrete species [45]. Note that this would need to be multiplied by the size of the genome measured in base pairs. We chose total genome mutation rates in the order of 10^0^ in our stochastic simulations, which is the estimated mutation rate for human somatic cells in various healthy tissues [36, 59]. However, our methodology holds for any realistic mutation rate and in particular for higher mutation rates.

### 2.2 Stochastic simulations

We use a Gillespie Algorithm [7] to implement our model in the Julia Programming Language [49]. Usually, stochastic simulations are in the order of 10^3^ - 10^6^ time units. If not otherwise stated, *t* = 10^5^ and *r* is set to 1. Every unit of time we record the VAF. We run 100 simulations for every parameter combination and analysis the patterns of mutation expansion and fixation. Every time unit can include multiple division events, as e.g. with a rate *r* = 1 on average *N* division events are included in one time unit. To avoid the dependence of our results to any artificially defined initial VAF distribution and to have resilience to small changes of population behaviour, we simulate an expansion phase from a single individual to the carrying capacity in all simulations. Thus, the birth-death constant population dynamics start with randomly accumulated mutations naturally evolving from a single individual.

## 3 Results

### 3.1 Joint fixation of multiple mutations

Changes of mutation frequencies are relatively slow in the Moran compared to the Wright-Fisher process, as only a single birth and death event are occurring at each time step. However, we still observe in stochastic simulations that multiple mutations fixate together, see for example figure 2, as opposed to a process of only sequential fixations of singular mutations. We denote the fixation of multiple mutations at the same time as a ‘jump’, and the number of mutations jointly fixating as the size of each ‘jump’. In a single simulation, the size of each jump varies and can at times reach extremely high values. These “jumps” can be observed even if looking at the number of fixed mutations over time averaged over multiple simulations, since the underlying exponential distribution of jump sizes tends to be dominated by single extremely large events.

First, we calculate the average number of mutations in a fixation event, i.e. jump size *J*_*N*_. We consider every fixation event from the point of view of the last mutation that turned up inside a jump. For example, if mutations *m*1, *m*2, *m*3, …, *m*_*l*_ fixate at the same time, the jump size of this fixation event is *l* and the last mutation in this jump is *m*_*l*_. Since it is the last mutation in a jump, we know that all mutations coming afterwards have not fixated at the moment of this ‘jump’. Similarly, all the mutations arising beforehand fixate in the population, if they were also present in the same individual where this ‘last’ mutation first appeared. If a mutation is present in *k* individuals, the chance for them to be present in any randomly chosen individual is *k/N*. Note that any mutation present in multiple individuals could fixate earlier instead of the jump we are currently looking at, but they are guaranteed to fixate eventually and that any mutation which is not present in this individual is guaranteed to to go extinct. Since these are all the mutations that could either fixate in the current ‘jump’ event or fixate earlier. This gives us an upper bound of the jump size. In addition, we know that the equilibrium VAF distribution of mutations is 2 *· Nµ/k* for a constant population [61]. As any fixation event also has a minimum size of 1, combining these observations we have

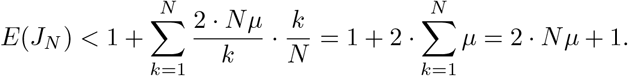

To make this estimate more precise, we look at the precise moment that the last mutation ‘*m*_*l*_’ in a jump just emerged in a newly born individual. The moment the offspring of this individual fixates, the jump happens. Investigating an arbitrarily chosen mutation ‘B’, which is present in this individual with a prevalence *k*, it is part of this jump only if it does not fixate beforehand. For this purpose, we can look at the sub-population composing of individuals without mutation *m*_*l*_, which has a population size of *N−* 1, and mutation ‘B’ has a prevalence of *k −* 1 in this sub-population. This implies a fixation probability of (*k −* 1)*/*(*N −* 1). We know that this sub-population is bound to extinction, but as there are no multiple deaths at once in our model, there will always be a time in which this sub-population only consists of a single individual. Since this last individual can either hold mutation ‘B’, in which case ‘B’ will have fixated in this sub-population, or not, in which case ‘B’ will have gone extinct, there is no possible ambiguity. If we now look at all mutations present in the same individual as mutation ‘*m*_*l*_’ instead of a single arbitrarily chosen mutation, we get the following estimate:

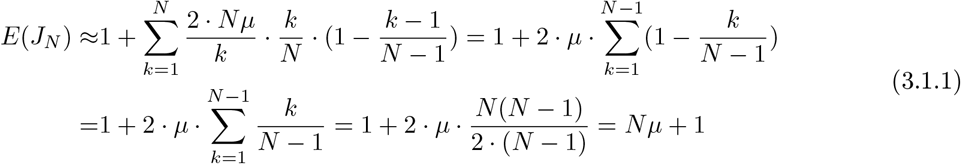

This implies that the average jump size grows linearly both with population size*N* as well as with mutation rate *µ*.

The estimate of the average jump size allows a direct estimate for the average ‘jump time’ *JT*_*N*_, which is the average time between two sequential jump events. We know that at every time step, on average 2*µ* mutations (*µ* per daughter cell) are generated, of which 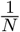 will fixate [5]. Using the estimate of how many mutations fixate at once (eq. 3.1.1), we get for the average jump time

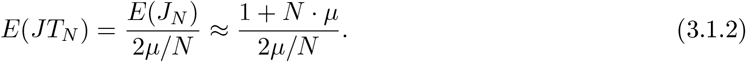

For 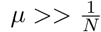, the 1 in *E*(*J*_*N*_) is negligible, which means

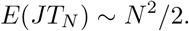

Note that this is counting every reproduction event as a time step. If you count time steps differently, for example generation time where every time step is *N* reproduction events as opposed to a single one, you speed up the dynamics by *N* and hence get 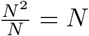 scaling in generation time.

For sufficiently high mutation rate *µ*, the average time between jumps grows with *N* ^2^ and is independent of *µ*. However, for the case of 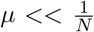, it grows linearly in *N* and is proportional to 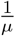. Interestingly, there is a transition in the scaling of the average jump time that depends on the population size *N* as well as the mutation rate *µ*. This implies that results can directly depend on the experimental setup, e.g. on the fraction of the genome considered for inference, as 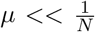 might hold true if looking at small parts of a big genome.

Note that accounting for a Poisson distributed influx of novel mutations is simple. By replacing the +1 in equation 3.1.1 and 3.1.2 with the appropriate mean of the Poisson distribution with a given *µ* excluding zero one gets the following equations:

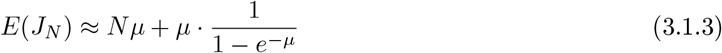

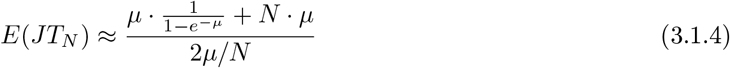

The general scaling behaviour of the two equations remains unchanged; For *µ <<* 1, the term 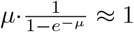, and for 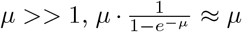. Around *µ ≈* 1, it never exceeds *≈* 2*µ*. In summary, the jump size linearly depends on both *N* and *µ*, while the jump time for ‘realistic’ values of *µ* depends only quadratically on *N*, but not on *µ*.

The jump time being independent of *µ* can be explained intuitively by looking at the fixation of offspring of individuals instead of the fixation of mutations. Necessarily for a mutation to fixate, the descendants of the first individual with that particular mutation need to take over the whole population. However, the converse is not true as there are “empty fixations”, in which the descendants of an individual fixate but they collectively hold no unique mutations that have not already fixated. Thus, no mutations fixate in this case. This becomes less and less likely with increasing *µ* and *N*, as correspondingly in the sub-population of descendants it is increasingly likely to have some unique mutations fixated already. Once *µ* and *N* are sufficiently high, a fixation of descendants of an individual without any fixation of corresponding unique mutations becomes extremely unlikely and as such the jump time is then independent of any further increase in *µ*. Figure 1 shows that our theoretical approximations agree well with our stochastic simulations under different parameters. Recall here that we count the time in the simulation in generations, i.e. we have on average *N* reproduction events every time step as opposed to to a single one. This does not change the fundamental dynamics in any way, but it means that the scaling of the jump time in generation time is linear instead of quadratic.

**Figure 1:**
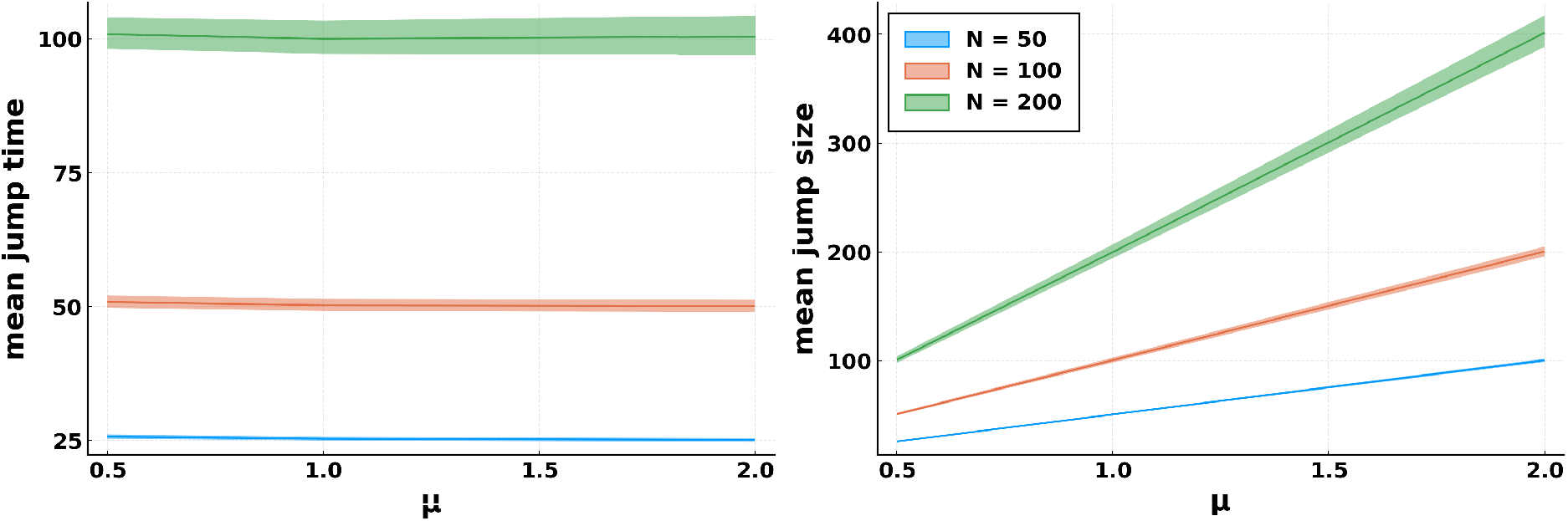
Mean jump time and jump size under different mutation rates *µ* and the population sizes *N*. In our stochastic simulations, the mean time between two sequential fixation events (jump time) is independent of mutation rate, but increases with the population size (left panel). The mean number of mutations in each fixation (jump size) increase with both the mutation rate and the population size. The green, red and blue area show the variance among simulations, and the solid lines refer to the mean values, which agree with our theoretical prediction given by equations 3.1.1 and 3.1.2 (100 simulations with 10^5^ time steps. Time in the simulations is measured in generations)

As expected from the theoretic expressions, while the time between two jumps appears to be independent of *µ*, the average size depends linearly on it. On the other hand, both values are linearly dependent on *N*.

Next we look at the distributions of jump sizes and jump times, going beyond the average values. The following argument gives us the correct distributions: the jump time *JT*_*N*_ is memoryless – after a fixation event the just-fixated mutations are “forgotten” and can not influence the time to the next fixation event. In addition, there is no mechanism making fixation more likely or less likely if there was no fixation for a while. The only memoryless discrete distribution is the geometric distribution, which can be directly projected into a continuous exponential distribution. The distribution of the jump size *J*_*N*_ on the other hand is on its own harder to estimate, but the two distributions are related, mediated by *N*. For *N* = 1, the distribution of *J*_*N*_ will be exactly the distribution of new mutations (usually a Poisson distribution), while *JT*_*N*_ will still be a geometric distribution dependent only on the likelihood of zero new mutations. A longer time between “fixations” will not have any impact on the size of the next fixation (the next jump size), meaning it will still be exactly the same Poisson distribution. On the other hand if *N* is sufficiently high, an increased time between fixations will on average lead to a linearly larger number of fixating mutations exactly as predicted, and result in jump size *J*_*N*_ also being exponentially distributed.

Our stochastic simulations confirm our rationale as shown in figure 2, the geometric distribution:

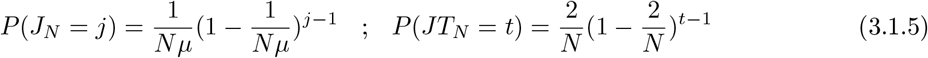

indeed fits very well for both *J*_*N*_ and *JT*_*N*_. Note that the exponential estimates in figure 2 are not fitted by our simulation data. Instead, we merely use the average value of jump time and jump size calculated before, and put them into geometric distributions, leaving no degree of freedom. In addition, the distribution of jump size implies that for a *N* >> 1, the chance for only a single mutation to fixate as opposed to multiple mutations fixating together is 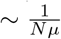.

**Figure 2:**
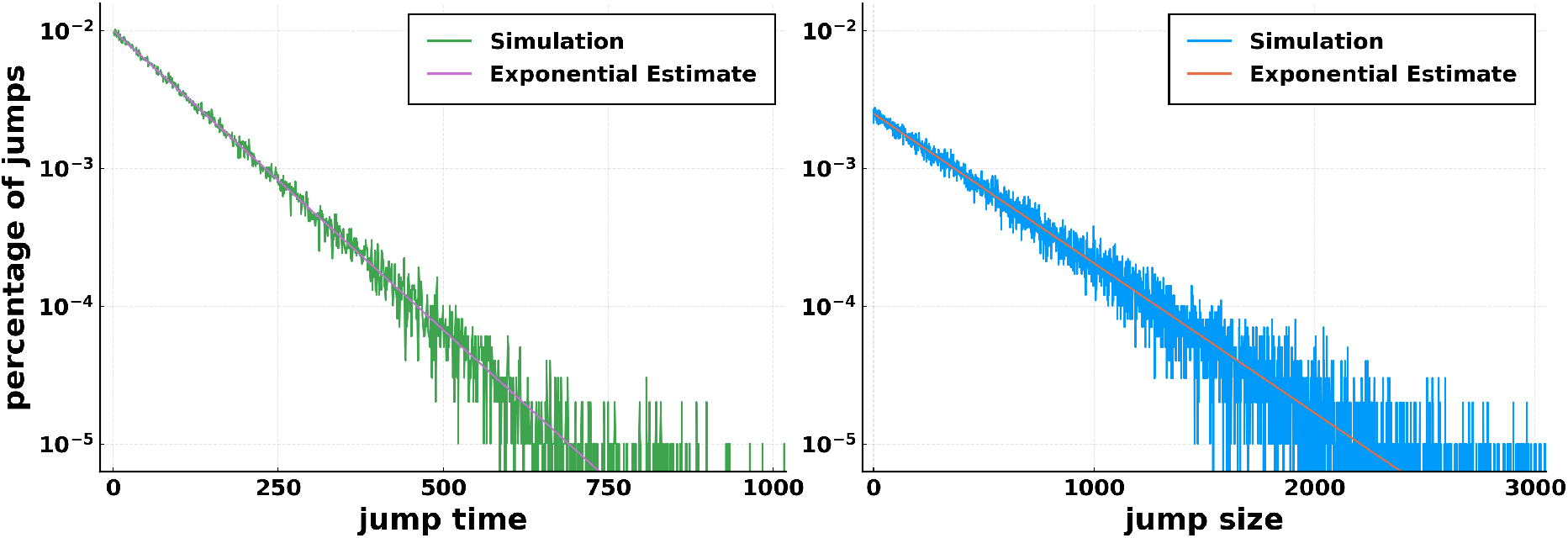
Distributions of jump time and jump size. Every jump refers to a fixation event of multiple mutations, the number of which is the jump size. We record all jumps as well as the time between two sequential jumps (jump time) over 100 simulations with 10^5^ time steps per simulation, based on which we obtain the distribution of jump time (green, left panel) and jump size (blue, right panel). We estimate the corresponding theoretical distribution based on the calculated average values (violet/red straight line). The geometric distribution is a very good fit for the data, and thus the theoretic average values appear to be correct (*N* = 200, *µ* = 2.0).

### 3.2 Accumulating wave distribution

The behaviour of “jumps” is not specific to the moment of fixation. It can also be observed when multiple mutations reach a high frequency simultaneously. This makes the term “jump” less fitting, and we introduce the concept of “accumulating waves”. Each wave is composed of multiple mutations. Especially at the higher frequencies, mutations tend to be strongly correlated up to being part of the same clone and as such move the frequencies up and down as one unitary wave, accumulating mutations over time until either fixation or, more likely, extinction. The “wave size” at any frequency is then defined as the number of mutations in each wave when measured while it is at that frequency.

Similar to the jumps, every instance of a “wave” is recorded as well as the corresponding number of mutations in each wave and the frequencies these waves pass by. Examples for the wave distributions under different mutation frequencies are shown in figure 3, where an exponential distribution is plotted against all wave sizes of a given frequency. While the wave size distribution at lowest frequencies bear little resemblance with the exponential distribution, those at high frequencies are well described by an exponential distribution, except for a heavier tail, culminating in an almost perfect fit absent variance around *f* > 0.9.

**Figure 3:**
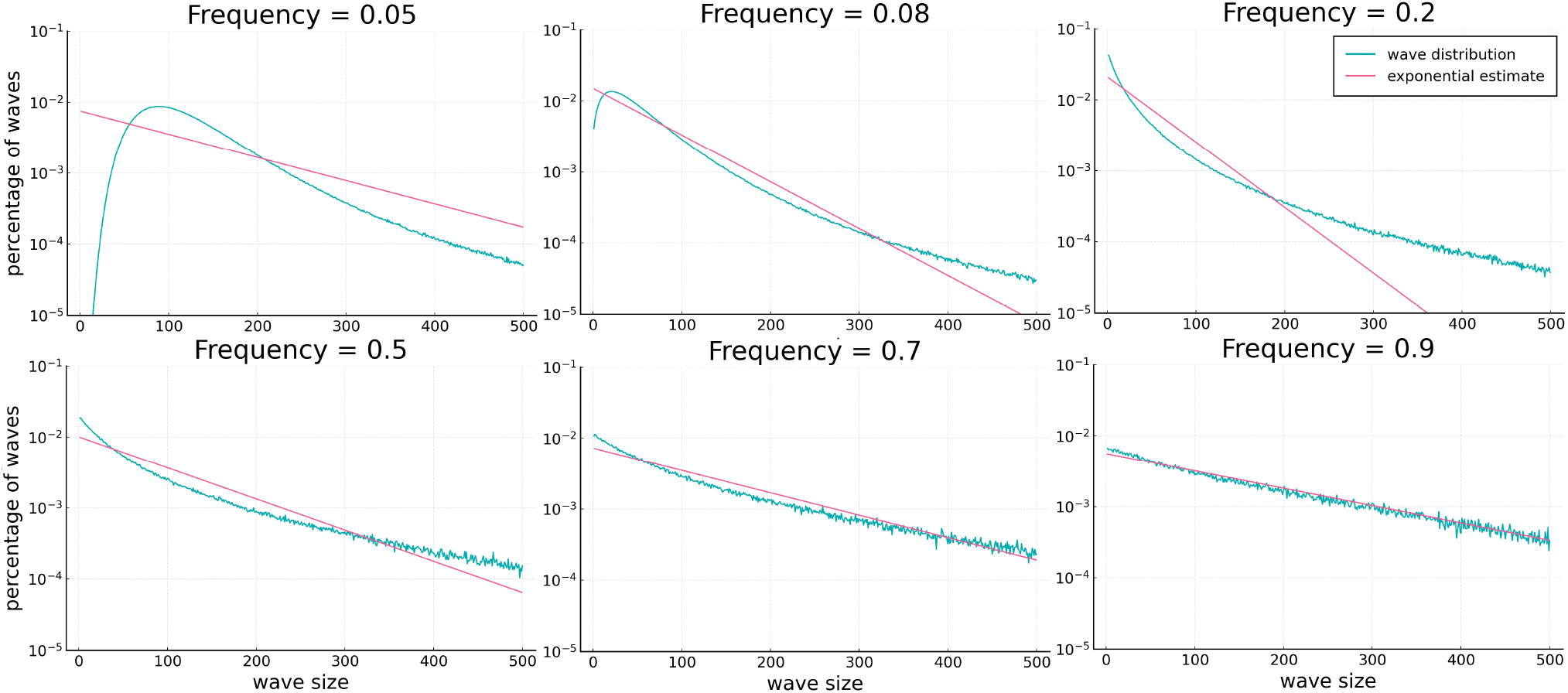
Distribution of wave sizes at different mutation frequencies. Mutations do not only fixation together, but also reach given frequencies at the same time, which we define as mutation “waves”. Each panel refers to a chosen mutation frequency, e.g. frequency = 0.9 refers to the distribution of mutation waves reaching 90% of the whole population. We show the percentage of waves (y-axis) as the number of waves carrying a given number of mutations (wave size, x-axis) divided by all waves in a simulation. At the higher frequencies, the distribution is very well approximated by an exponential distribution. At intermediate frequencies (for example 0.5), the distribution has a considerably heavier tail than the exponential, i.e. very high wave sizes are much more likely than the exponential predicts. At low frequencies (for example 0.05), the difference is even more pronounced and the distribution has a clear intermediate maximum, while the exponential is continuously falling (averaged over 100 simulations, 10^5^ time steps per simulation, N = 100, *µ* = 2.0).

A possible explanation is the fundamentally different wave dynamics at different mutation frequencies. As already described in section 3.1, fixations which is equivalent to mutation frequency at 1, are dominated entirely by waves that have accumulated from their inception as a single individual. This results in an exponential distribution. Mid-to high frequencies are largely dominated by similar waves. Those mutations jointly reaching mid- and high frequencies are more likely to be carried by the same subgroup of individuals, or by clustering subgroups. However, since there is still a small number of waves that have reached even higher in frequency before falling back to the current frequency, the wave distribution has a longer tail. In contrast, mutation waves at low frequencies are dominated by the influx of new mutations with minimal accumulation over time. Those mutations jointly reach the same value of low frequencies are usually present in multiple distinct groups of individuals, all of which add up to generate the VAF at that frequency. Since this is mostly the sum of largely independent Poisson distributions, the result approaches a normal distribution. As there are still substantial numbers of waves moving from higher to lower frequencies, often with large numbers of “locked” mutations, i.e. those mutations are no present in independent subgroups of individuals, the wave distribution is still significantly skewed and has a much longer tail compared to a regular normal distribution. An approximation is shown in supplementary figure 9.

We quantify the fitting of the wave distribution with its corresponding exponential estimates as “exponential fit index”, which is the absolute distance between the wave size distribution and the corresponding fitted exponential distribution for all mutation frequencies and under various population sizes (see figure 4). Absent the discontinuity that is present in a single frequency around *f* = 0.5, the fit is consistently getting better for the higher frequencies. A slight leveling of the fit can be observed at the highest frequencies, which are true for all population sizes but more obvious for larger population sizes at *N* = 200. This leveling is caused by mutation waves reaching a very high frequency are less compared to the case in relatively lower frequencies, and thus have a higher variance. Since the mutation waves need more time to reach the very highest frequencies when the population size increases, this leveling is more obvious for larger populations under the same simulation time. In a summary, the increasingly good fit of the exponential distribution with the wave distributions shows that the concept of accumulating waves dominating the dynamics of large part of the VAF is far more general than the original concept of jumps which only apply to the moment of fixation.

**Figure 4:**
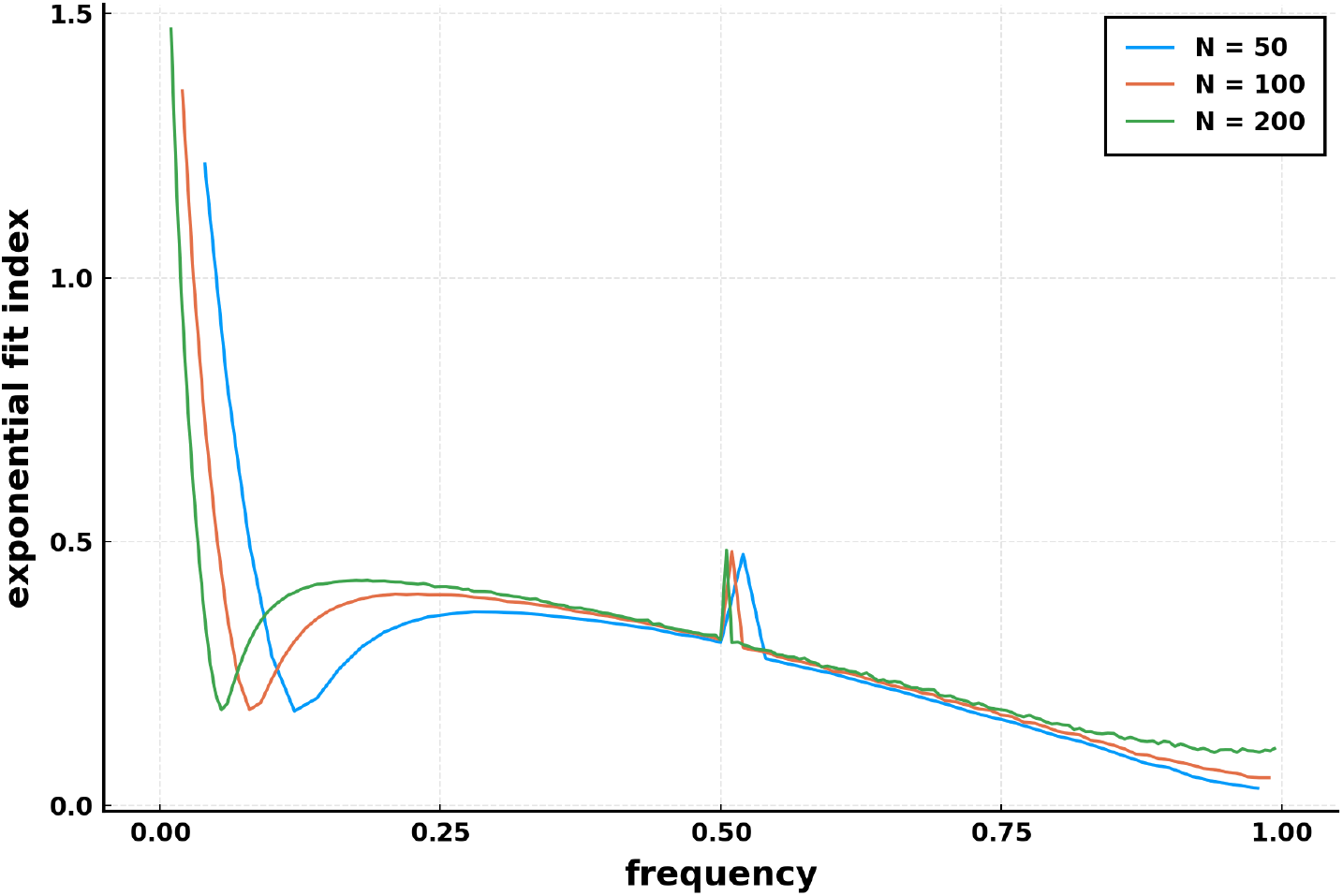
Statistics of the fitting between the wave size distribution and the exponential distribution for different frequencies and population sizes *N*. We obtain the “exponential fit index”, based on the absolute distance between the wave size distribution in our stochastic simulations and its corresponding fitted exponential distribution. The squared distance or distances in the log scale give similar results. If fit index = 0, the distribution is identical to the exponential distribution. While the very low frequencies are clearly far from the exponential distribution, the fit becomes much better when frequency increases (around frequency = 0.2 for *N* = 50 and frequency= 0.1 for *N* = 200) and are very good for high frequencies.(100 simulations, 10^5^ time steps per simulation, *µ* = 2.0)

### 3.3 Average wave frequency distribution

To understand how these waves are related to the variant allele frequency, we introduce the wave frequency distribution (WFD). It represents the average number of mutations in waves (average wave size) reaching a given frequency. The WFD is very similar to the VAF, which measures the number of mutations having a given frequency over time. The difference is that in the WFD we have to exclude all time points where there are no waves, which means at those time points no mutations are present at given frequencies. We calculated WFD as the mean of all recorded waves over the entire time span of each simulation, and present the average as well as the variance cross multiple simulations. Figure 5 shows the WFD for *N* = 100, *µ* = 1.0. The shape of the WFD is the same for all other populations *N* and *µ* but varies in absolute values (See figure 7). Since the lower frequencies are never empty and always have mutations, the WFD is identical to the VAF there. However, for the higher frequencies the WFD significantly deviates from the VAF, as it measures the size of the mutation waves passing by. Those waves can either move further to fixation, back down towards extinction, or just fluctuating around. This shows that for single evolutionary repeats there is a bi-modal nature of the variant allele frequencies. For most of the time, there are no mutations present in many especially high frequencies, but occasionally there is a growing wave of mutations passing by. Instead, a continuous VAF distribution with a power law decay over frequencies is an average over time or over multiple evolutionary repeats under constant neutral populations. For a single evolutionary process measured at a given time point, a VAF distribution exceeding the expected average by large margins is not necessarily an aberration but due to the bi-model nature of VAF.

**Figure 5:**
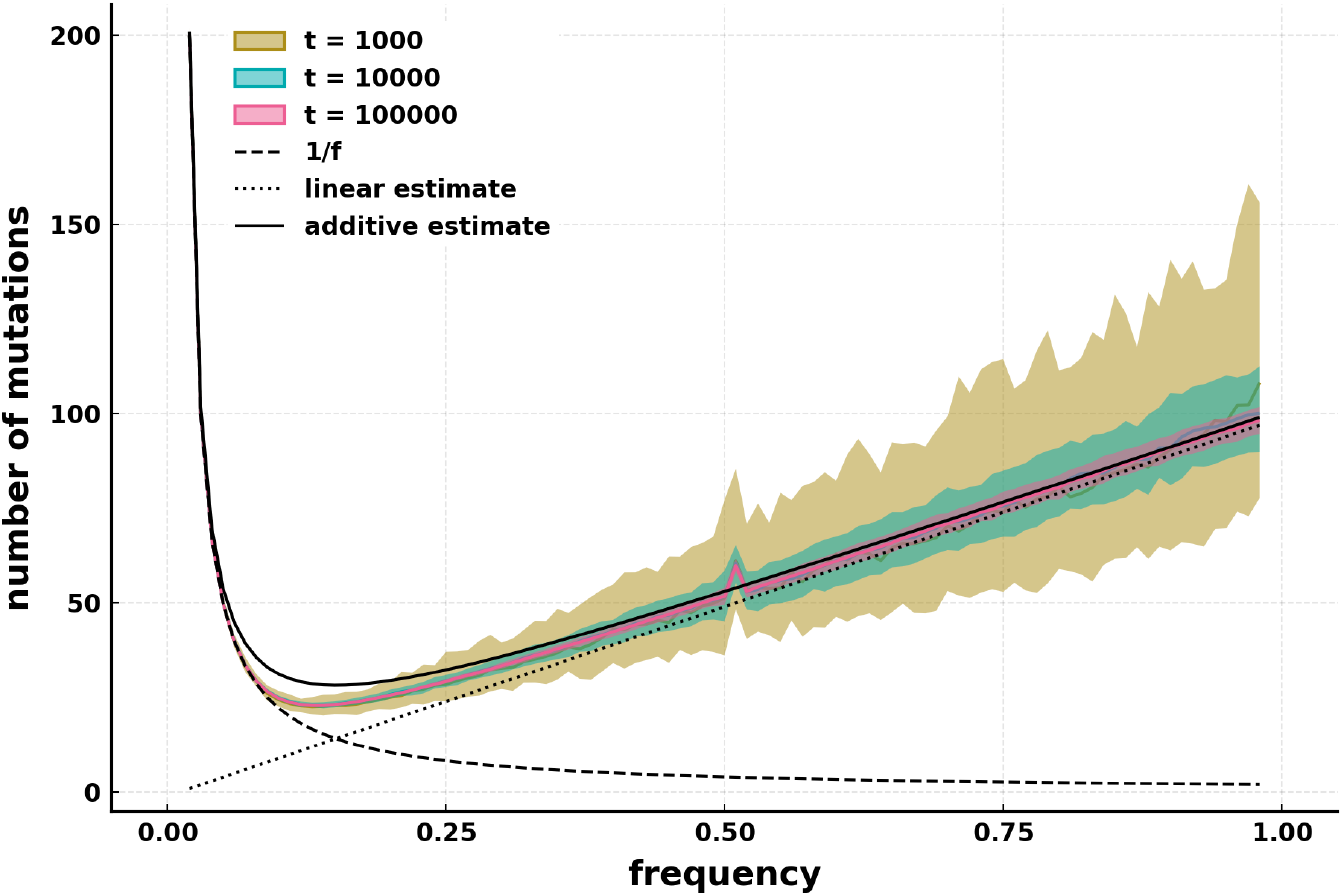
Average wave sizes against mutation frequencies. Shown is the average number of mutations reaching a given frequency at the same time (wave size) and the standard deviation (as a colored area) from multiple stochastic simulations. The wave size at each frequency is the mean of all measured waves at that frequency over the entire time span of each simulation, i.e. it is the mean of the distribution shown in figure 3. We compare this distribution with the expected variant allele frequency distribution (VAF, dashed line), i.e. a 1*/f* shape for constant populations, as well as the prediction from the wave size estimate (dotted line) and an upper limit (solid line) by simply adding both the VAF and the wave size estimate. For the lower frequencies, the two distributions are almost identical. For example at frequency one, new mutations are emerging every single time step in new individuals, so all the mutations simultaneously becoming “locked” with one another into a single wave never happens. The expected VAF distribution and the measured wave distribution deviate from each other when frequency increases, just as the likelihood of singular waves increases. Roughly from frequency = 0.2 onward, the wave sizes progresses linearly as predicted from the wave size estimation. We show that the distribution of the mean number of mutations reaching each frequencies is stable and consistent over time (*t* = 1000, 10000, 10000), but the variance decreases (100 simulations, 10^5^ time steps per simulation, *N* = 100, *µ* = 1).

If we assume that each frequency is its own sub-population and reaching that frequency is akin to fixating, we can use equation 3.1.3 to estimate the average WFD

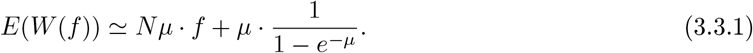

Interestingly, although this assumption is only reasonable for the very highest frequencies, the resulting estimate fits well even for frequencies as low as *f* = 0.25. This shows again how the estimates we derive from the very specific concept of a jump/wave are not only applicable for high frequency where only one wave can be present at a time, but actually even for frequencies where, in principle, two waves are possible. For example at frequency 0.25, we can in principle have up to 4 different sub-populations, each at frequency 0.25, with their respective fixated mutations constituting 4 different waves. But since that frequency is on average largely empty of mutations, it’s in practice rare for multiple waves to interfere, and hence it largely behaves as if only one wave was possible.

#### 3.3.1 Estimating the time without mutations per frequency

Following our explanation in the above section, we can check at which frequencies the WFD and VAF differs by investigating the amount of time there are no mutations at a given frequency, *T* (*f*). Our model not only implies that many frequencies are empty with no mutations for most of time, but it also gives a specific estimate. We know that the average number of variants for frequency *f* over time and/or multiple simulations is *V* (*f*) *≃* 2*µ/f*, the average wave size for the same frequency is 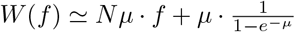, and *V* (*f*) = (1 *− T* (*f*))*W* (*f*) + *T* (*f*) *\** 0 due to the bi-model nature of VAF. Thus, the fraction of time no mutations are present at a given frequency *f* is approximately

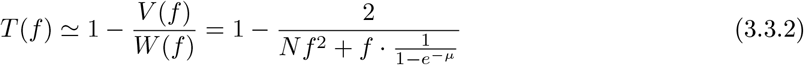

This theoretical estimate agrees well with “true” values recorded directly in our stochastic simulations (see Figure 6). As can be seen, for small N the estimate is already very good for *f ∼* 0.5 and higher, while for larger N the estimate is almost indistinguishable from the true value from *f ∼* 0.25 onward. *V* (*f*) >> *W* (*f*) implies the presence of multiple waves, which is true for *∼ f <* 0.2 for *N* = 50 and *∼ f <* 0.1 for N = 200. Our WFD estimate is generally more conservative in this respect than the value derived from simulations, meaning it serves as a lower bound for the time a frequency is empty of mutations.

**Figure 6:**
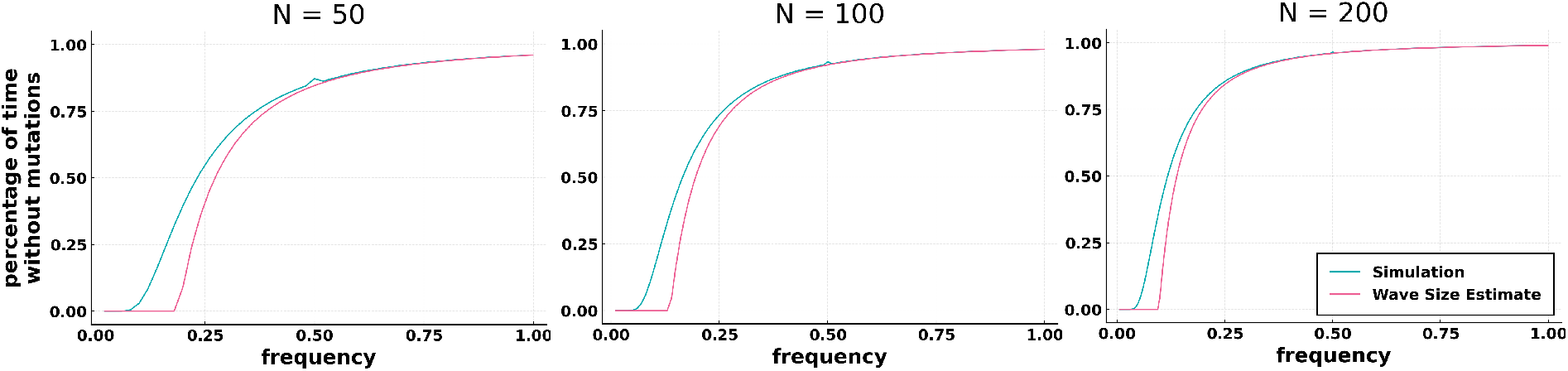
Percentage of time a given frequency has no mutations present on the y-Axis, the respective frequency on the x-Axis. For example, at the lower frequencies up to ca 0.05, the VAF values will almost never be zero in the simulation, while close to 1.0, it will be zero for most of the time. For the higher frequencies, we can predict this with only a small error by simply estimating the average wave size and dividing the average VAF at that point by the average wave size. (100 simulations, 10^5^ time steps per simulation, *µ* = 1)

## 4 Discussion

Simultaneous fixation of multiple mutations have been reported in many experimental data sets [12, 14, 27, 44, 61]. This can be explained by accumulation through mutators, clonal interference or negative frequency dependent selection of beneficial mutations [14, 38, 39, 42, 44, 47], but can also happen in neutral dynamics. We have quantified and analysed the joint fixations of random neutral mutations and proposed a new concept of accumulating mutation waves in constant populations.

In the context of the rise of cheaper and more extensive genome sequencing, analyzing the variant allele frequency distribution of natural populations in experiments has become much more feasible. Likewise, the theoretic expected distribution has been explored for a number of different models already [37, 48, 55, 58, 60, 61]. However, we show that in addition to the traditional approach of looking only at the average numbers of mutations at each frequency over time, we can explain the distribution in much closer detail by separating out the time when each frequency are empty with zero mutation and the time when mutations are present. We show how the specific distribution at each frequency follows an exponential tail that grows larger with increasing frequency, until encompassing the entire distribution at fixation. This results in the concept of “accumulating waves”, which means that for large parts of the frequency spectrum, the dynamics are dominated by mutation waves that are on average far larger than expected from an averaged VAF.

The predictions for the average wave size at each frequency stemming from our wave size estimate describe results from stochastic simulations well. For reasonable population sizes (*N ≥* 50) at frequencies *∼ f ≥*0.5, the average simulations are almost indistinguishable from the linear estimate. The entire distribution are within bounds defined by the power law VAF distribution, the linear estimate, and the sum of both even after varying population parameters (see Appendix A). Using the relation between the traditional VAF estimate *V* (*f*) and our WFD estimate *W* (*f*), we derive an estimate for the expected time a frequency is entirely empty of mutations, which fits reasonably well for *f* > 0.25 and are almost indistinguishable for *f* > 0.5 compared to our stochastic simulations. Correspondingly, in any particular realization of a simulation, the recorded non-zero VAF values in higher frequencies usually far exceed the expectation by traditional power law VAF estimates in constant neutral populations. We can again reliably predict the average magnitude of these values, since they on average follow our linear WFD estimate. There are some concepts related to the wave frequency distribution, e.g. the clone size distributions, the spatial size of a given clone [46], has been used to calculate compound population statistics.

Overall, our approach extends the concept of the VAF and not only explains the notable phenomena of large numbers of simultaneous mutations fixating in a very simple model without requiring either selection or more specific population dynamics like negative frequency dependent selection, but also directly gives a prediction for the shape of the distributions as well as the average expected values. It shows that the VAF distribution is functionally very different for high and low mutation frequencies, due to the presence of multiple mutation waves at the lower frequencies and single waves at the higher frequencies. For large parts of the frequency spectrum especially for high frequencies, the VAF is effectively bimodal – empty for the most part, but waves that can get extremely large are occasionally passing by. Lower frequencies are instead split up into a quasi-normal distribution for small wave size, and an exponential tail that follows the exponential predicted from the jump size distribution.

This approach can in principle be used to further analyze population dynamics of experimental populations, but most practical applications would require even more extensive data than we have available. As more and better genetic data becomes available, we will increasingly be able to analyze it with more information-rich and detailed concepts, instead of being relegated to simpler to measure but also less informative concepts.

## Acknowledgements

BW is supported by a Barts Charity Lectureship (grant MGU045), as well as an UKRI Future Leaders Fellowship.

## Appendices

### A Varying population size N, mutation rate *µ* and time *t*

To control how robust our model is we already varied some parameters in the preceding sections and figures. In figure 7, an overview over the effects of varying either *N* or *µ* is shown. Each figure is in the same style as figure 5, meaning it shows the wave sizes on the y-Axis against the frequencies on the x-Axis. Again, each data point is the average of 100 simulations. In general, varying each value has different effects.

**Figure 7:**
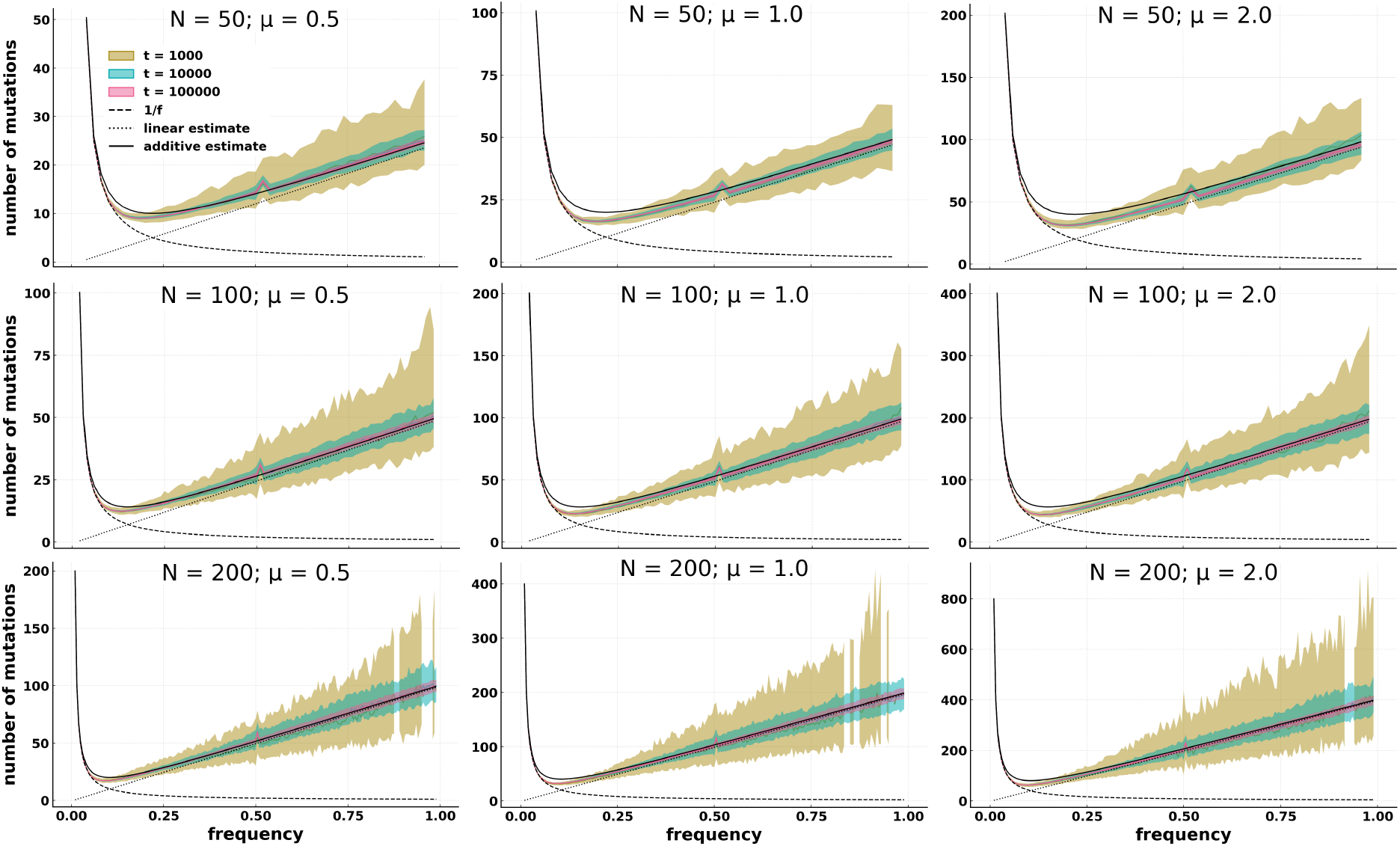
Average wave size on the y-Axis against a given frequency on the x-Axis. Even after varying the population size and mutation rate, the accumulating wave linear estimate is still a very good fit for the high frequency distribution. From top to bottom, increasing population size. From left to right, increasing mutation rate. The empty spots for *t* = 1000 and *N* = 200 are due to not having enough data points to calculate a variance. However, for *t* = 10000 and higher, all frequencies are sufficiently represented.

Similar to figure 5, increasing the time *t* does not substantively change the distribution for any combination of N an *µ*, but the increased number of data points reduces the variance. In other words, after *∼* 1000 division events per cell, the equilibrium state has already been all but reached for the chosen values.

Increasing the population size *N*, on the other hand, does change the distribution notably. As can be seen, the higher *N*, the larger the part of the distribution that fits the linear accumulating wave estimate, as opposed to the 1*/f* VAF estimate. In addition, both the lowest frequencies as well as the very highest frequencies increase linearly in amplitude with *N*, e.g. for *µ* = 0.5 the lowest frequency is *W* (1*/N*) *∼ N*, while the highest frequency is *W* ((*N −* 1)*/N*) *∼ N/*2. Only the lowest point of the distribution, around *f ∼* 0.1, does not change in amplitude nearly as much as the aforementioned frequencies, it only moves slightly to the left in terms of frequency, overall making the distribution much steeper for increasing *N*. At last, increasing the mutation rate *µ* simply increases the entire distribution linearly, but changes nothing in the shape of the distribution.

In summary the changes from varying either *N* or *µ* in the higher frequencies fit very well with the predictions from 3.1.1 for the chosen values. Similarly, the changes in the lower portion of the distribution also fit expectations considering that for higher N, the 1*/f* becomes steeper independent of the wave size distribution and hence the intersection between the 1*/f* and the linear *f* necessarily has to happen earlier.

Notably, no matter how the values are chosen, the average of the WFD at all frequencies never seems to fall below the lower limit given by either the 1*/f* or the linear *f* estimate.

### B A local peak at *f* = 0.5

In the figures 4, 5, 6 and 7, a “bump” can be seen at the frequency right above *f* = 0.5 for dynamics under different population sizes, i.e. *f* = 0.52 for *N* = 50, *f* = 0.51 for *N* = 100 and *f* = 0.505 for *N* = 200. In clonal reproduction as in our model, partially overlapping mutations are impossible. Any two randomly chosen mutations A and B have to either exclude one another, where individuals carrying A and B belong to two different sub-populations, or include one another where the sub-population carrying one mutation must include all individuals carrying the other mutation. That means a discontinuity happens at *f* = 0.5 - up to this value, two (or more) distinct waves with the same frequencies are possible, while beyond it, only one wave is possible. That means the very next value after *f* = 0.5 is part of a fundamentally different distribution. While this is not a direct explanation how this local peak phenomenon in the wave frequency distribution comes to be, it is a possible hint in the right direction.

We can also show it in the same way as the exponential distributions (Figure 8), where we compare the frequency *f* = 0.51 to its direct neighbours, *f* = 0.5 and *f* = 0.52. Despite these close neighbours usually being very similar in shape, *f* = 0.51 deviates much more strongly than the other two from the exponential.

**Figure 8:**
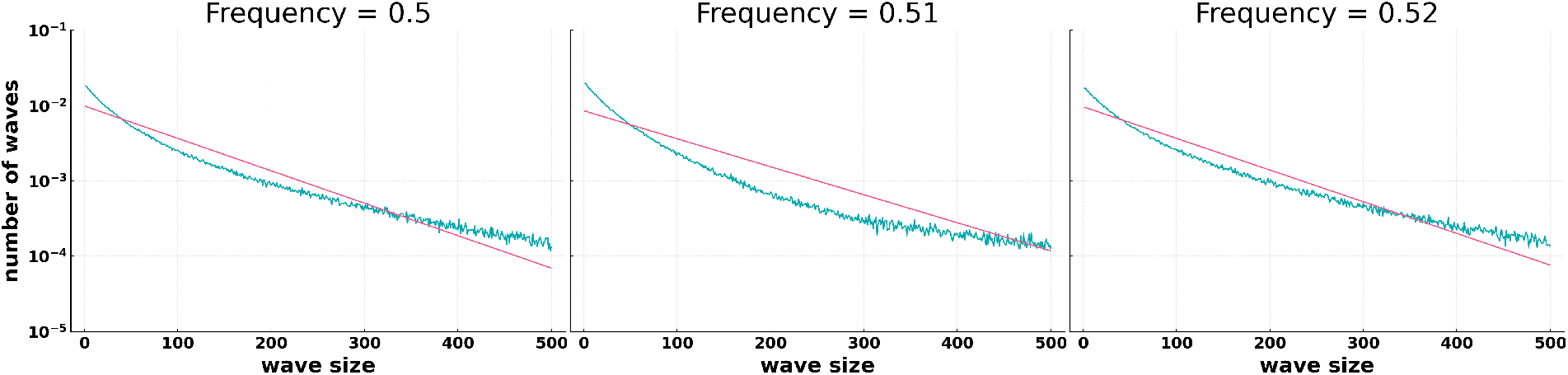
Example of the anomaly around *f* = 0.5 for *N* = 100 and *µ* = 1. The relative number of waves on the y-Axis against the wave size on the x-Axis. While *f* = 0.5 and *f* = 0.52 are already quite close to the expected exponential distribution, *f* = 0.51 is much more skewed and deviates relatively strongly from the exponential

### C Low Frequency Wave Distribution

**Figure 9:**
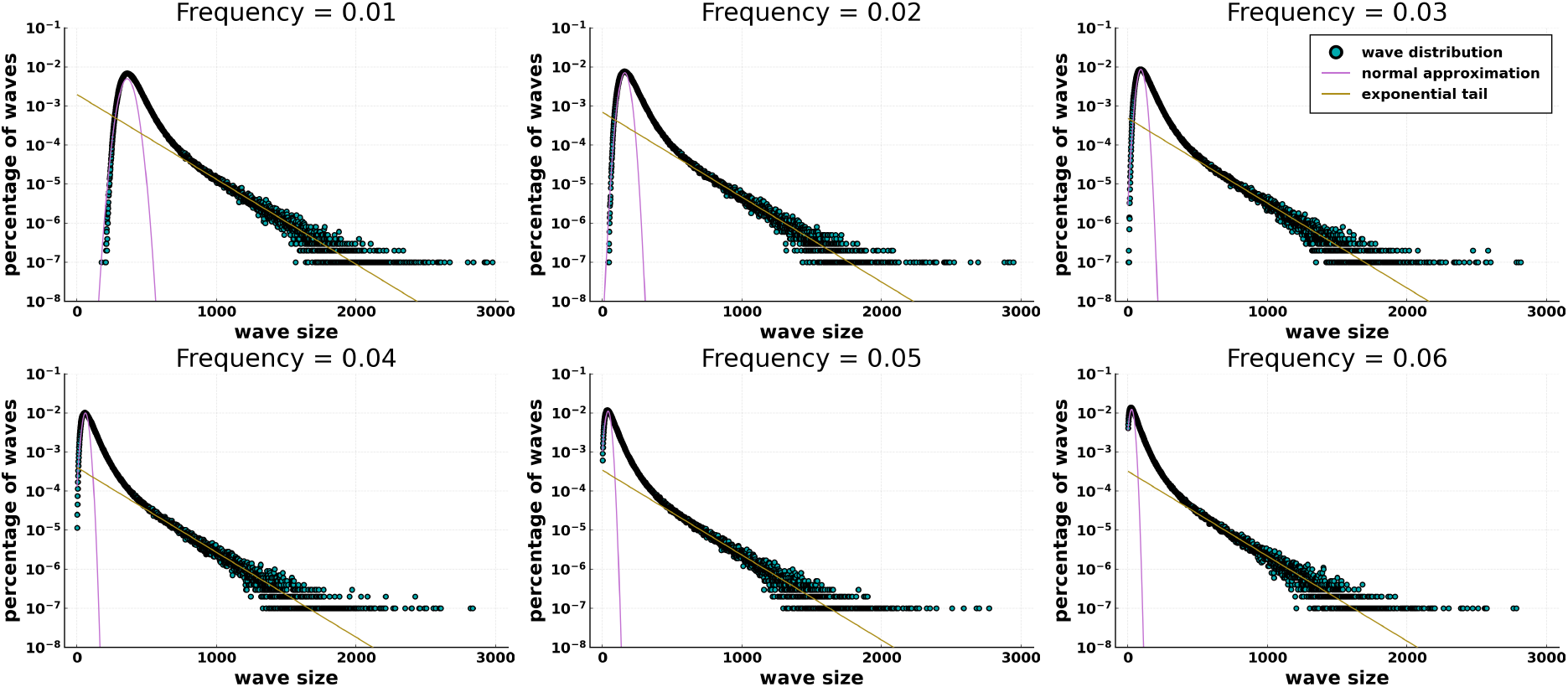
Distribution of waves sizes at different mutation frequencies. Each panel refers to a chosen mutation frequency, e.g. Frequency = 0.01 refers to the distribution of mutation waves reaching 1% of the whole population. We show the percentage of waves (y-axis) as the number of waves carrying a given number of mutations (wave size, x-axis) divided by all waves in a simulation. This figure specifically shows the 6 lowest frequencies in a population of size *N* = 100. Unlike the higher frequencies, these frequencies have a skewed normal distribution as the bulk of their distribution, but they have a non-normal exponential tail that follows exactly the prediction from the jump size distribution in terms of slope

